# Assessment of Neurovascular Coupling & Cortical Spreading Depression in Mixed Models of Atherosclerosis & Alzheimer’s Disease

**DOI:** 10.1101/2020.08.13.249987

**Authors:** Osman Shabir, Ben Pendry, Llywelyn Lee, Beth Eyre, Paul Sharp, Monica A Rebollar, Clare Howarth, Paul R Heath, Stephen B Wharton, Sheila E Francis, Jason Berwick

## Abstract

Neurovascular coupling is a critical brain mechanism whereby changes to blood flow accompany localised neural activity. The breakdown of neurovascular coupling is linked to the development and progression of several neurological conditions including dementia. In this study, we examined cortical haemodynamics in preparations that modelled Alzheimer’s disease (J20-AD) and atherosclerosis (PCSK9-ATH) between 9-12m of age. We report novel findings with atherosclerosis where neurovascular decline is characterised by significantly reduced blood volume, levels of oxyhaemoglobin & deoxyhaemoglobin, in addition to global neuroinflammation. In the comorbid mixed model (J20-PCSK9-MIX), we report a 3x fold increase in hippocampal amyloid-beta plaques. A key finding was that cortical spreading depression (CSD) due to electrode insertion into the brain was worse in the diseased animals and led to a prolonged period of hypoxia. These findings suggest that systemic atherosclerosis can be detrimental to neurovascular health and that having cardiovascular comorbidities can exacerbate pre-existing Alzheimer’s-related amyloid-plaques.

## Introduction

Alzheimer’s disease (AD) is the most common form of dementia worldwide, with the vast majority of cases being sporadic and occurring 65 years and over. Population based studies have shown that AD and vascular pathologies commonly coexist in the brains of elderly individuals (Kapasi et al., 2017; Matthews et al., 2009; Neuropathology Group. Medical Research Council Cognitive & Aging, 2001; Rahimi & Kovacs, 2014). A major cardiovascular pathology that affects as many as up to 60% of all individuals after the age of 55 is atherosclerosis. Atherosclerosis is the progressive thickening, hardening and narrowing of major arteries, including those that supply the brain, such as the carotids (Lusis, 2000). Intracranial atherosclerosis does not occur until much later in life, around 75 years and above. As such, Alzheimer’s disease that begins around the 8^th^ decade of life is usually present with other comorbidities such as atherosclerosis (Napoli et al., 1999). There is also evidence that, not only do these often exist as comorbidities, but they may interact pathogenically with vascular disease and neurovascular unit changes contributing to AD (Iadecola, 2017; Kapasi & Schneider, 2016). To date, there are very limited models of comorbidity with respect to preclinical studies, and instead models have been very specific and ‘pure’, and not reflective of the clinical pathology in humans. Atherosclerosis is known to be a major risk factor for the development of dementia. The progressive atheromatous plaque build-up within cerebral arteries that supply the cortex over time can lead to stenosis producing insufficient oxygen delivery to the brain parenchyma, potentially resulting in neuronal death and symptoms of dementia (Shabir et al., 2018). Indeed, the vascular cognitive impairment (VCI) which precedes the onset of dementia may be attributed to a variety of different vascular pathologies affecting either systemic or intracranial vasculature (both large or small vessels) (Iadecola et al., 2019). Due to the complexity of atherosclerosis and dementia pathogenesis, understanding the mechanisms of their mutual interactions is necessary if efforts to develop therapeutics to prevent VCI and vascular dementia, which currently has no disease-modifying cure, are to succeed.

The breakdown of NVC is thought to be an important and early pathogenic mechanism in the onset and progression of a range of neurological conditions (Zlokovic, 2011). In the present study, we aimed to investigate neurovascular function in mid-aged (9-12m old) mice where atherosclerosis was a comorbidity. We used a novel model of atherosclerosis that utilises a single adeno-associated virus (AAV) i.v. injection of a gain of function mutation (D377Y) to proprotein convertase subtilisin/kexin type 9 (rAAV8-mPCSK9-D377Y), combined with a high-fat Western diet to induce atherosclerosis in most adult mouse strains (Bjorklund et al., 2014; Roche-Molina et al., 2015). This leads to the constitutively active inhibition of the LDL-receptor preventing cholesterol internalisation and degradation by hepatocytes, leading to hypercholesterolaemia to occur and the development of robust atherosclerotic lesions within 6-8 weeks (Bjorklund et al., 2014). Furthermore, in order to address the effect atherosclerosis could have on mild Alzheimer’s pathology, we combined the atherosclerosis with the mild J20-hAPP mouse model of familial Alzheimer’s disease (fAD) to create a mixed comorbid mouse model (J20-PCSK9-MIX). The J20-hAPP mouse model of fAD over-expresses human amyloid precursor protein (hAPP) with the Swedish (K670N and M671L) and the Indiana (V7171F) familial mutations (Mucke et al., 2000), which begin to develop amyloid-beta (Aβ) plaques around 5-6 months of age, and show signs of cognitive impairments from 4 months (Ameen-Ali et al., 2019). We hypothesised that atherosclerosis would exacerbate Alzheimer’s disease pathology in the brain and that neurovascular function would be further worsened compared to AD or ATH models alone. We have previously reported no significant alterations to evoked-haemodynamics in the J20-AD model of the same age (9-12m); however, under acute imaging sessions where an electrode was inserted into the brain, we found significantly perturbed haemodynamics (Sharp et al., 2019). We hypothesised that electrode insertion causes cortical spreading depression (CSD). Based on recent data linking migraine with aura with cardiovascular disease (Kurth et al., 2020), we hypothesised that experimental CSD might be heightened in all disease models. We report that experimentally induced atherosclerosis in the J20-AD model increased the number of Aβ plaques by 300%. Furthermore, experimental CSD is more severe in all diseased groups compared to WT controls.

## Results

### 2D-Optical Imaging Spectroscopy (2D-OIS) Measures Brain Cortical Haemodynamics Through a Thinned Cranial Window

We performed chronic imaging of the brain cortex 3-weeks post-surgery, where the thinned cranial window remained intact (Figure 1A/B), as described previously (Shabir et al., 2020; Sharp et al., 2019). We deployed a range of stimulations (2s & 16s mechanical whisker stimulations) with the mouse breathing both 100% oxygen (hyperoxia) and 21% oxygen (normoxia), in addition to recording transitions between conditions and performing a 10% hypercapnia test to test the maximum dilation of vessels. Each experimental day consisted of the same set of experiments with consistent timings to ensure reliability across all animal groups. First, a 2s-whisker stimulation (5Hz) with the mouse breathing 100% oxygen; hyperoxia, consisting of 30 trials, second, a 16s-whisker stimulation consisting of 15 trials. Animals were then transitioned from hyperoxia to 21% oxygen; normoxia, and the baseline haemodynamic changes were recorded. The same set of stimulations were deployed under normoxia (2s & 16s stimulations), before transitioning back to hyperoxia for a final 10% hypercapnia test. Using these stimulations, activation maps of blood volume; total haemoglobin (HbT), can be generated (Figure 1C). Mice were allowed to recover and after 1-week, a final acute imaging session was performed. In this setup, a small burr-hole was drilled through the thinned skull overlying the active region of interest (ROI) as determined from the chronic imaging sessions (Figure 1D), and a multichannel electrode was inserted into the brain (Figure 1E) to record neural activity simultaneously. We then imaged and recorded the baseline haemodynamics for a 35-minute period to observe the effect electrode insertion, before commencing the first stimulation. This was also done to record baselines on chronic imaging sessions.

**Figure 1:**
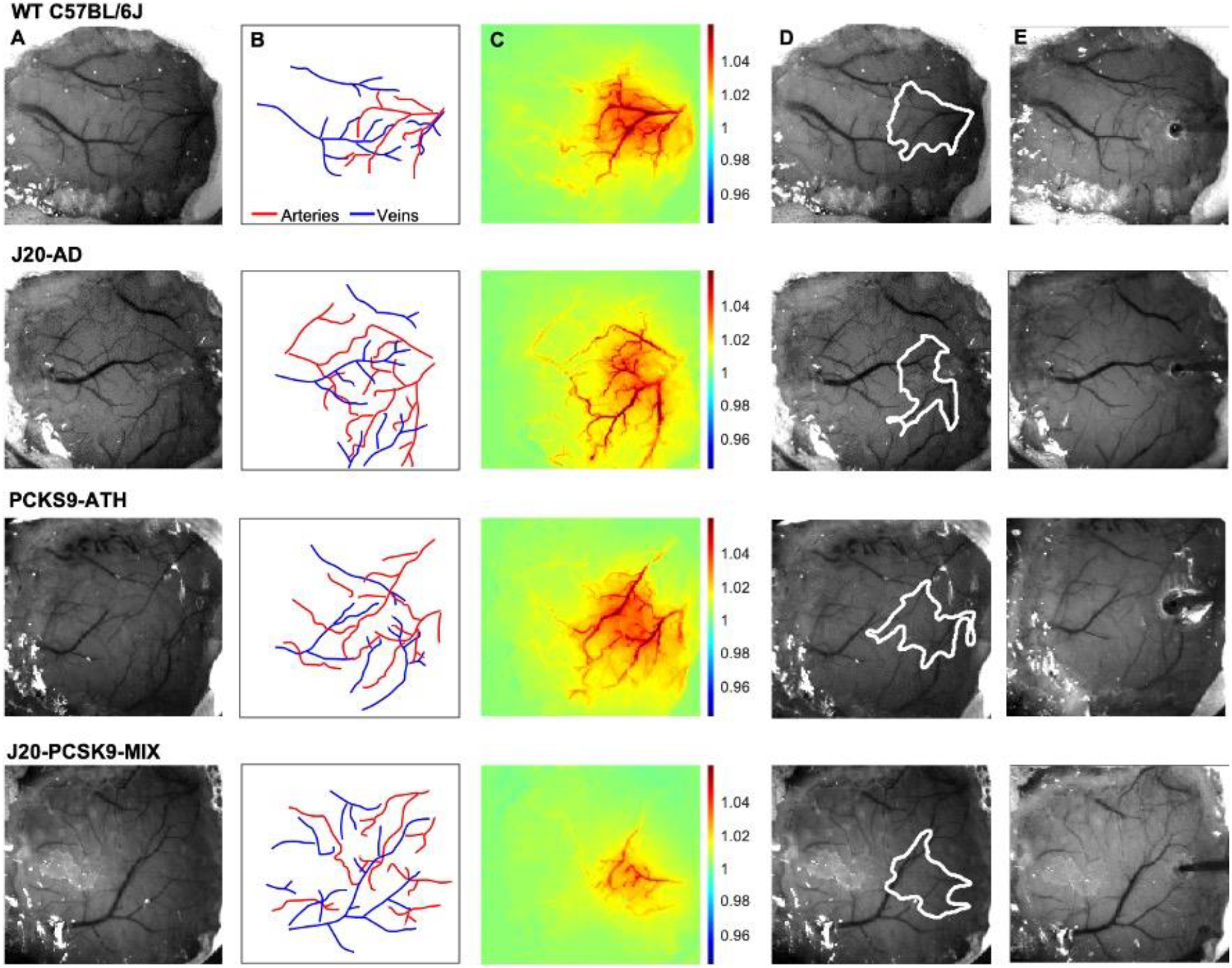
Experimental Setup and Data Derivation. **A)** Raw image of representative thinned cranial windows for WT, J20-AD, PCKS9-ATH AND J20-PCSK9-MIX mice (chronic imaging session). **B)** Vessel map outlining the major arteries and veins within the thinned cranial window. **C)** HbT spatial activation map showing fractional changes in HbT in response to a 16s-whisker stimulation. **D)** Automated computer-generated region of interest (ROI) determined from the HbT activation response in C from which time-series for HbT, HbO & HbR are generated. **E)** Raw image of the same animals in terminal acute imaging sessions with multichannel electrodes inserted into the active ROI determined from chronic imaging session.

### Chronic Haemodynamic Responses in the Brain are Reduced in PCSK9-ATH Mice

Cortical haemodynamics were imaged through a thinned cranial window to determine whether evoked cortical haemodynamics were different between 9-12m old wild-type (WT), atherosclerotic (PCSK9-ATH), Alzheimer’s (J20-AD) & mixed (J20-PCSK9-MIX) mouse models (Figure 2). Across all stimulations and conditions, ATH-PCSK9 mice displayed a significant reduction of evoked blood volume responses (HbT; peak value) compared to WT controls. J20-AD mice and J20-PCSK9-MIX mice did not exhibit a significant change in HbT across all stimulation conditions compared to WT mice. Evoked HbT responses; although initially are smaller in J20-PCSK9-MIX mice, recovered to match that of J20-AD mice later in the experimental protocol under normoxia (Figure 2D). Levels of oxyhaemoglobin (HbO) were significantly reduced in PCSK9-ATH mice but showed a reduced trend in J20-PCSK9-MIX mice too. The washout of deoxyhaemoglobin (HbR) was significantly reduced in PCSK9-ATH mice compared to WT, but also showed a reduced trend across all diseased groups across all conditions compared to WT mice. All mice displayed stable and robust haemodynamic responses across the experimental protocol (Figure S2). Finally, vascular reactivity as determined by the response to 10% hypercapnia was not significantly different between any of the diseased groups (Figure S3).

**Figure 2:**
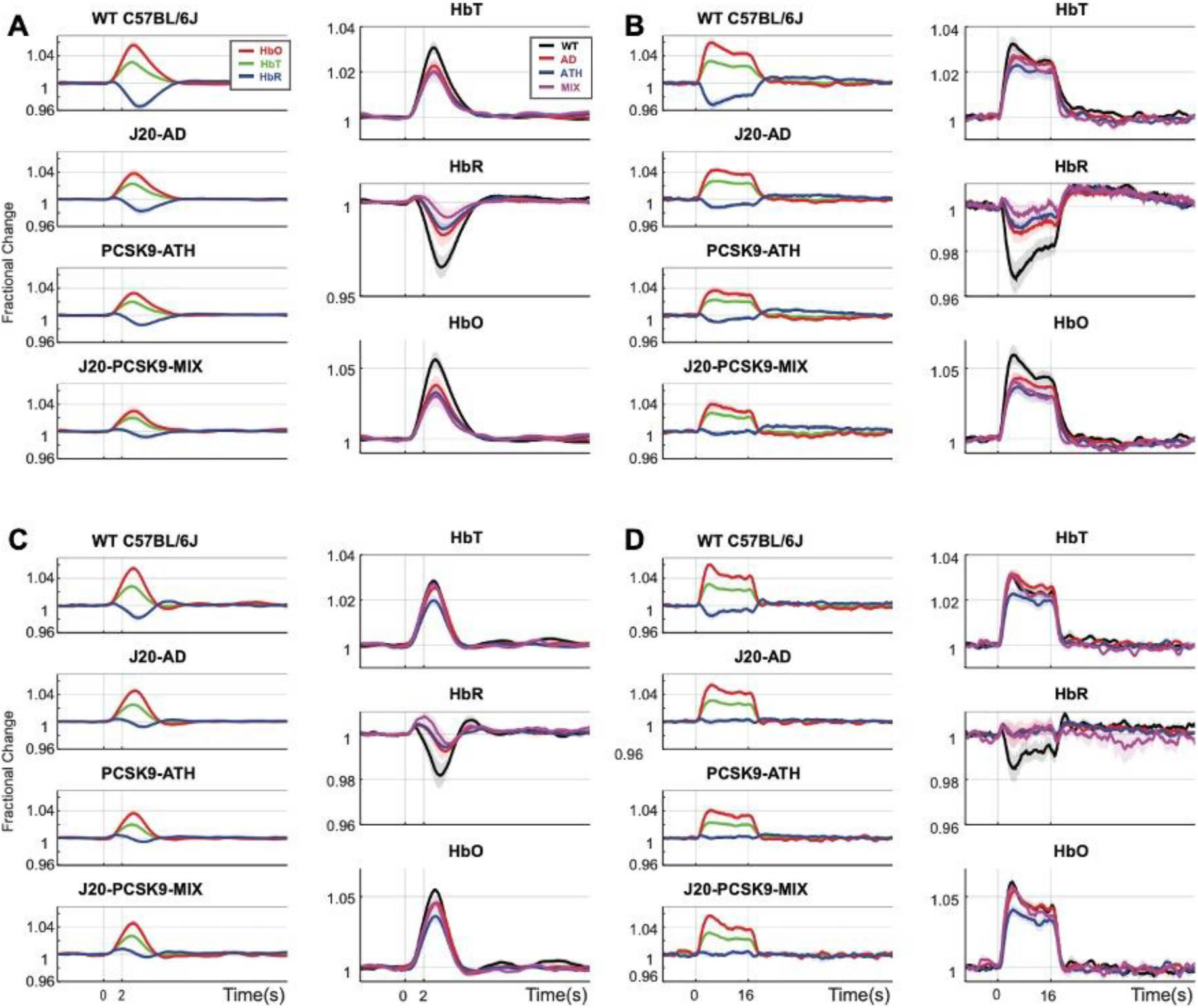
Fractional Changes in Chronic Stimulus-Evoked (Peak) Haemodynamic Responses. **A)** 2s-stimulation in 100% oxygen. **B)**16-stimulation in 100% oxygen. **C)**2s-stimulation in 21% oxygen. **D)**16s-stimulation in 21% oxygen. All animals aged 9-12m: WT (n=6), J20-AD (n=9), PCSK9-ATH (n=8), J20-PCSK9-MIX (n=6). **HbT:** There was no significant overall effect of disease F(3,25)=2.83, p=0.059. However, Dunnett’s (two-sided) multiple comparisons test revealed there was a significant difference between WT and ATH (p=0.023). As expected, there was a significant effect of experiment, F(1.65,41.14)=13.64, p<0.001. There was also no significant interaction effect between experiment and disease, F(4.94,41.14)=1.50, p=0.211. **HbO**: There was a significant overall effect of disease F(3,25)=4.84, p=0.009. Dunnett’s (two-sided) multiple comparisons test revealed there was a significant difference between WT and ATH (p= 0.002). There was a significant effect of experiment, F(1.47,36.72)=15.348, p<0.001. There was no significant interaction effect between experiment and disease, F(4.41,36.72)=1.64, p=0.181. **HbR**: There was a significant overall effect of disease F(3,25)=4.86, p=0.008. Games-Howell multiple comparisons reveal HbR peak is significantly different for WT vs ATH (p=0.040). There was a significant effect of experiment, F(1.69,42.28)=17.33, p<0.001. There was a significant interaction between experiment and disease interaction: F(5.07, 42.28)=3.19, p=0.015. All error bars (lightly shaded) are ±SEM. Vertical dotted lines indicate start and end of stimulations.

### CSD is Worse in Diseased Animals and Impacts Haemodynamic Recovery to Baseline

1-week after recovery from the chronic imaging protocol, an acute imaging experiment was performed wherein a small-burr hole was drilled into the skull overlying the active region (determined from HbT responses from chronic experiments) and a microelectrode was inserted into the brain to a depth of 1500-1600μm to obtain neural electrophysiology data in combination with the imaging of cortical haemodynamics by 2D-OIS. Electrode insertion into the brain resulted in a wave of haemodynamic changes that occurred in all mice (CSD) (Figure 3). In WT mice, electrode insertion led to a small decrease in HbT (vasoconstriction) followed by a robust HbT bounce back (vasodilation), immediately followed by a small, sustained vasoconstriction (reduced HbT) that persisted for some time (Figure 3A-top). In J20-AD mice, electrode insertion caused a large vasoconstriction to occur which spread across the cortex in a strong wave of vasoconstriction that was followed by a very small, attempted recovery. This was masked by a large sustained and prolonged vasoconstriction of contiguous vessels that persisted for some time (Figure 3A-bottom). The largest vasoconstriction post-CSD occurred in J20-AD mice, followed by PCSK9-ATH mice, then J20-PCSK9-MIX mice compared to WT controls. The smallest of all CSD occurred in WT mice (Figure 3B). A prolonged and sustained constriction below baseline persisted in all mice post-CSD, however, this effect was recovered to baseline in WT mice during the first stimulation experiment 35 minutes after the CSD occurred (Figure 3B). In all disease mouse models, the constriction below baseline was more severe and persisted for a much longer time with a slower haemodynamic recovery. Following CSD, stimulation-evoked haemodynamic changes were not significantly different in any of the diseased groups overall, although they were initially smaller in the first two stimulations for PCSK9-ATH mice (Figure S1).

**Figure 3:**
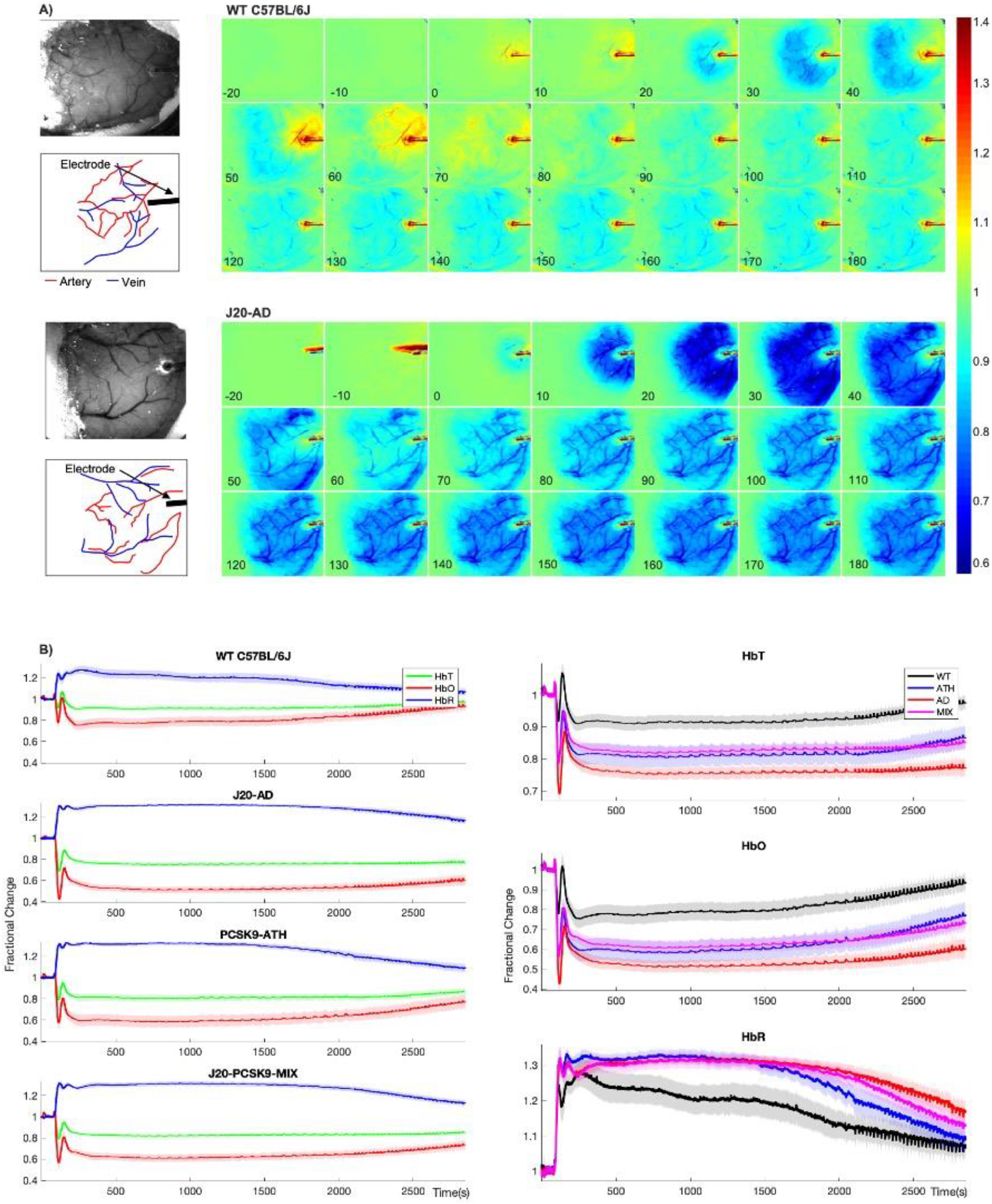
Cortical Spreading Depression (CSD) in WT, diseased and comorbid animals. **A)** Representative montage time-series of WT and J20-AD mice showing HbT changes post-electrode insertion. Colour bar represents fractional changes in HbT from baseline. **B)***Left:* Average CSD haemodynamics profiles for control animals (WT C57BL/6J & nNOS-ChR2) (n=7), J20-AD (n=7), PCSK9-ATH (n=5) & J20-PCSK9-MIX (n=6) mice. *Right:* Averaged changes to HbT (top), HbO (middle) & HbR (bottom) upon CSD in the different mouse groups compared to WT. **HbT:** A 1-way ANOVA showed significant effect of disease for HbT (F(3,21)=9.62, p=0.001. Dunnett’s 2-sided multiple comparisons showed that AD vs WT p<0.001, ATH vs WT p=0.012 & MIX vs WT p=0.020. **HbO:**1-way ANOVA showed significant effect of disease for HbO (F(3,21)=8.51, p<0.001. Dunnett’s 2-sided multiple comparisons showed that AD vs WT p<0.001, ATH vs WT p=0.01 & MIX vs WT p=0.017. **HbR:** Kruskal-Wallis test revealed no significant effect of disease H(3)=6.58, p=0.087. All error bars (lightly shaded) are ±SEM.

### Stimulus-Evoked Neural Activity is Not Significantly Altered in Any Disease Groups Compared to WT Mice

In the final imaging session and after a 35-minute period of recovery post-electrode insertion, the first experimental stimulation was performed (2s-stimulation in 100% oxygen) where evoked cortical haemodynamics were imaged simultaneously with the recording of neural multi-unit activity (MUA). Evoked-MUA response were not significantly different in any of the diseased groups compared to WT mice (Figure 4), suggesting that the significantly different evoked-HbT in PCSK9-ATH mice (observed on chronic imaging sessions) was due to neurovascular breakdown. Initially, the MUA was slightly lower for J20-AD, PCSK9-ATH & J20-PCSK9-MIX mice compared to WT mice (Figure 4A), however, later in the experimental session by the last stimulation, there was no observable difference in MUA between any of the groups (Figure 4D). Thus, this suggests that the neural MUA was initially smaller after the CSD had occurred, however, recovered fully with time. The haemodynamic responses in the acute experimental session were not significantly different across all stimulations for any of the diseased groups.

**Figure 4:**
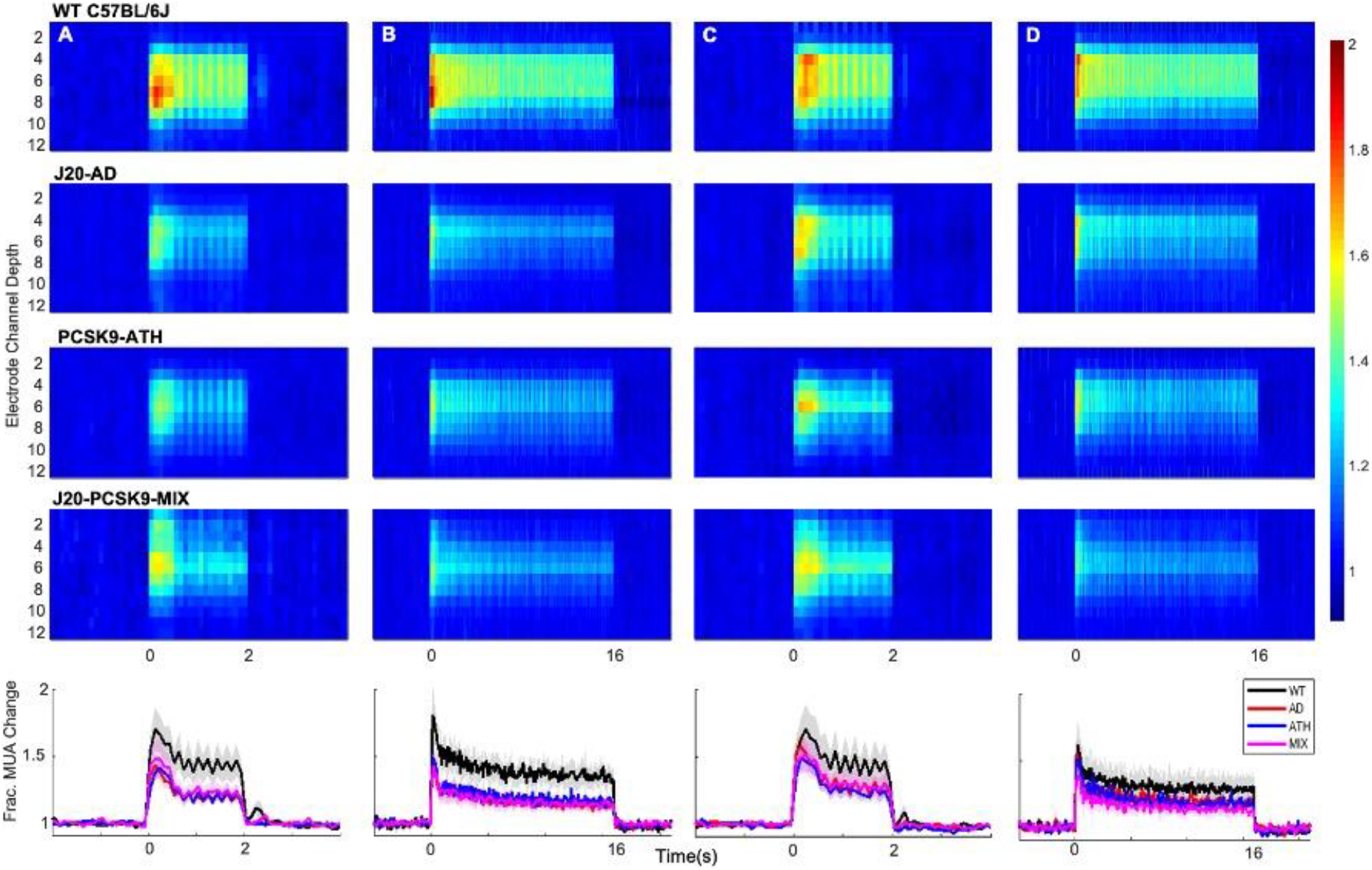
Evoked Neural Multi-Unit Activity (MUA) Responses. MUA heat maps showing fractional changes in MUA along the depth of the cortex (channels 4-8) in response to stimulations in WT C57BL/6J (n=6), J20-AD (n=9), PCSK9-ATH (n=7) & J20-PCSK9-MIX (n=6) mice. Overall effect of disease on MUA F(3,24)=2.24, p=0.109 (2-way mixed design ANOVA). There was a significant effect of experiment, as expected, F(2.26, 54.16)=6.83, p=0.002. There was no significant interaction between experiment and disease F(6.77, 54.16)=0.70, p=0.670. All error bars (lightly shaded) are ±SEM.

### Increased Number of Hippocampal Aβ Plaques in J20-PCSK9-MIX Mice. Increased Neuroinflammation in J20-AD and PCSK9-ATH Mice

Immunohistochemistry was performed on J20-AD and J20-PCSJK9-MIX mice to assess whether there were any specific differences in AD neuropathology changes. Staining was performed for Aβ plaques and these were quantified within the hippocampus and the cortex. Aβ plaques were significantly increased by 3-fold in the hippocampi of J20-PCSK9-MIX mice compared to J20-AD mice (Figure 5A/B). Within the cortex, there was no significant difference in Aβ plaques between the 2 groups (data not shown). Next, neuroinflammation was assessed by qRT-PCR for 2 key inflammatory markers: interleukin-1β (IL1β) and tumour necrosis factor-α (TNFα) to assess the degree of neuroinflammation present globally within the brain. IL1β mRNA was significantly upregulated in J20-AD and PCSK9-ATH mice (Figure 5C). TNFα mRNA was significantly upregulated in PCSK9-ATH mice only (Figure 5D). J20-PCSK9-MIX mice displayed the lowest inflammatory changes in IL1β & TNFα compared to the other diseased groups, though this was still higher than WT mice.

**Figure 5:**
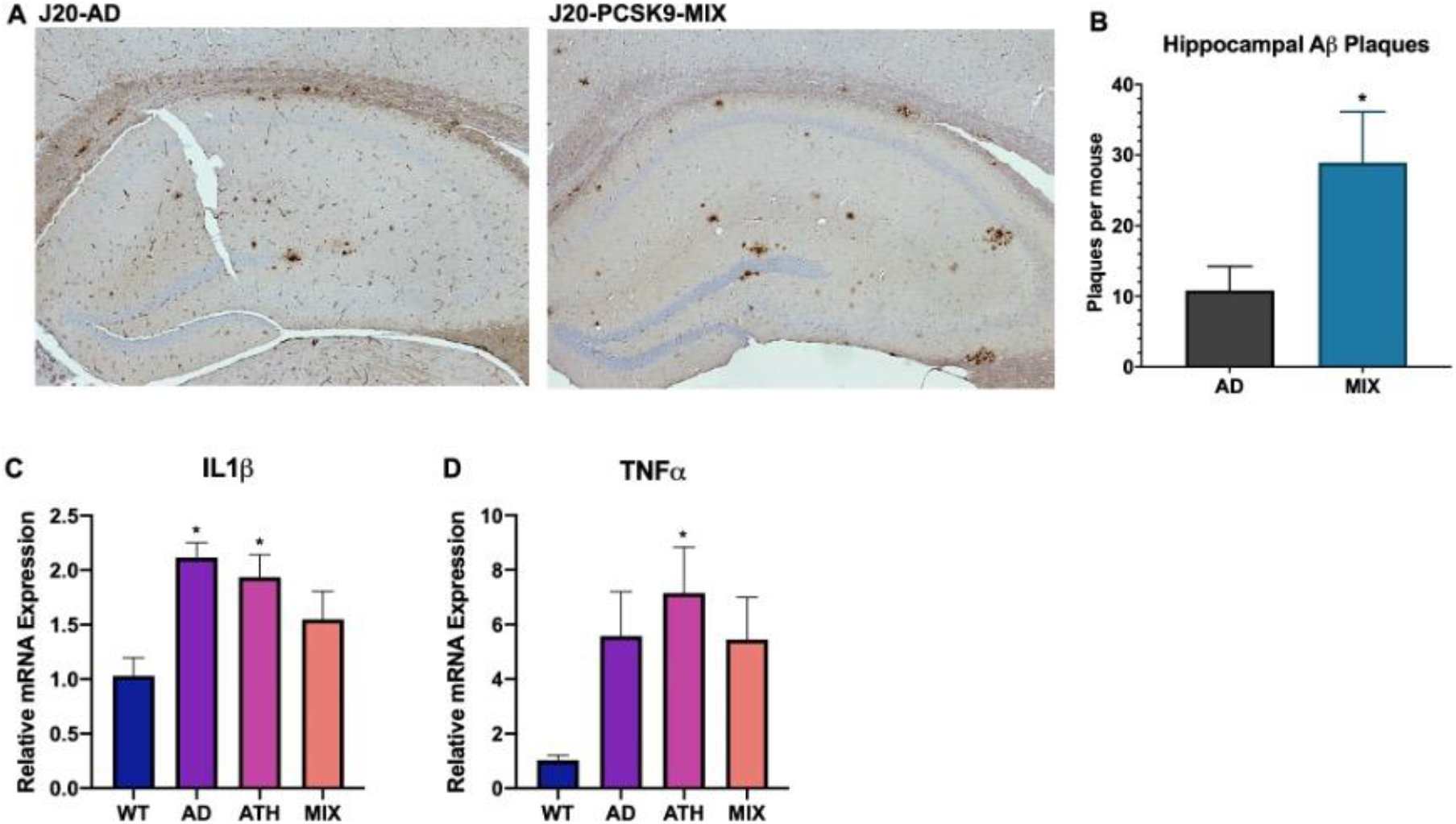
Neuropathology and Neuroinflammation. A) Representative histological coronal hippocampal sections for J20-AD and J20-PCSK9-MIX mice stained with anti-Aβ to visualise Aβ plaques. B) Increased number of amyloid-beta plaques in the hippocampus of J20-PCSK9-MIX mice compared to J20-AD mice (p=0.036; unpaired t-test) (n=4 each). Cortical plaques p=0.3372 (data not shown). D) qRT-PCR for 1L1β: AD vs WT p=0.011, ATH vs WT p=0.0278, MIX vs WT p=0.218 (1-way ANOVA with post-hoc Dunnett’s multiple comparisons test). E) qRT-PCR for TNFα: AD vs WT p=0.1197, ATH vs WT p=0.0370, MIX vs WT p=0.1313 with post-hoc Dunnett’s multiple comparisons test. All error bars are ±SEM.

## Discussion

The present study investigated neurovascular function in a novel experimental model of atherosclerosis (PCSK9-ATH) and for the first time, in a comorbid setting whereby atherosclerosis was experimentally induced in a well characterised model of AD; J20-hAPP, to create a mixed comorbid model (J20-PCSK9-MIX). These mice were compared to age-matched (9-12m) WT C57BL/6J controls, and J20-AD mice. Given that systemic atherosclerosis is a major risk factor for dementia, the mechanisms underpinning the relationship between atherosclerosis, neurovascular decline and dementia are still largely unclear.

In the first part of the study, we characterised evoked-haemodynamic responses using a chronic skull-intact & surgery-recovered mouse preparation. We found that PCSK9-ATH mice displayed significantly reduced evoked blood volume (HbT) responses, in addition to reduced levels of oxyhaemoglobin (HbO) and notably, an impaired washout of deoxyhaemoglobin (HbR) across all stimulations and conditions. The J20-PCSK9-MIX mice did not display a significant reduction in HbT, nor in HbO or HbR levels. With respect to J20-AD mice, we did not observe any significant alterations to HbT as previously published (Sharp et al., 2019). Another important finding from the present study was that 10%-hypercapnia responses were not significantly different in any of the mice compared to WT controls (Figure S3), thus suggesting that vascular reactivity was not impaired in any of the mice, indicating that cerebral arterioles were unaffected by atherosclerosis at this time-point (9-12m). Thus, the basis of reduced HbT in PCSK9-ATH mice cannot be attributed to intracranial atherosclerosis. A recent study corroborated our findings showing that cerebrovascular reactivity and cerebral blood flow was preserved in the Tg-SwDI model of Alzheimer’s (Munting et al., 2021), similar to our J20-model.

In the second part of the study, we obtained neural multi-unit activity (MUA) by inserting a multichannel electrode into the active region defined from the chronic imaging experiments. As we showed in our previous reports (Shabir et al., 2020; Sharp et al., 2019), the technical procedure of electrode insertion causes a cortical spreading depression (CSD) to occur in all animals. Here, we describe the CSD and its recovery on the different disease groups. CSD has two distinct phases: 1) a wave of depolarisation within the grey matter characterised by neuronal distortion leading to a large change of the membrane potential whereby neuronal activity is silenced (spreading depression) & 2) haemodynamic changes that accompany neuronal spreading depolarisation which typically result in a wave of prolonged reduced perfusion that persists for some time (Ayata & Lauritzen, 2015; Dreier, 2011). CSD does not typically occur in healthy brain tissue, however, it is a common neurophysiological occurrence in certain pathological conditions including migraine, epilepsy, brain injury, hyperthermia, hypoxia & ischaemia (Dreier, 2011). In WT mice, the initial constriction wave is small, and a robust haemodynamic recovery occurs which allows for neurovascular coupling to occur to sustain neurons metabolically. This is a marked difference to the diseased animals, which upon electrode insertion to cause a CSD, exhibit profound vasoconstriction with an extremely limited haemodynamic recovery resulting in a prolonged constriction, severe reductions to blood volume and HbO & HbR levels indicating hypoxia and ischaemia. Thus, the haemodynamic response to CSD in diseased mice is severely inappropriate and can lead to long lasting devastating effects such as widespread cortical pannecrosis of neurons and astrocytes (Dreier, 2011). As our data show, baseline blood volumes do not recover in the diseased animals for a much longer period compared to WT animals, with the most profound CSD occurring in J20-AD mice, followed by PCSK9-ATH mice.

CSD may be the neuropathological link between migraine, stroke, cardiovascular disease and dementia in which cardiovascular risk factors, genetics and other lifestyle factors which prime the onset of migraine to occur lead to vascular vulnerability within the brain predisposing affected individuals to an increased risk of cerebral ischaemia and haemorrhagic stroke (Ripa et al., 2015). There is accumulating evidence to suggest that shared genetic and associated clinical features observed in migraine patients are involved in the increased vulnerability to cerebral ischaemia, therefore, predisposing affected individuals to stroke and white matter lesions associated with dementia (Yemisci & Eikermann-Haerter, 2019). The underlying mechanism being CSD; the neurophysiological feature of aura in migraines, whose induction threshold can be reduced by genetic mutations and systemic comorbidities that contribute to vascular dysfunction and neuroinflammation (Yemisci & Eikermann-Haerter, 2019). Indeed, mouse models of cerebral autosomal dominant arteriopathy with subcortical infarcts and leukoencephalopathy syndrome (CADASIL); a genetic cerebrovascular disease caused by *NOTCH3* mutations that has a high frequency of migraines with aura, have enhanced CSD linking a dysfunctional neurovascular unit with migraine with aura (Eikermann-Haerter et al., 2011). Furthermore, a recent study examined women who suffered from migraines with and without aura and found that those that suffered migraines with aura had a higher incidence rate of cardiovascular disease compared to women without aura or any migraines (Kurth et al., 2020). In addition, another recent study found that migraine history was positively associated with an increased risk of developing both all-cause dementia and AD, but not VaD (Morton et al., 2019). Our study, along with the previously discussed studies provide an explanation for these recent findings and highlights how systemic disease can prime the brain to allow profound CSDs to occur in the context of migraine, and as such, migraine frequency and intensity may be related to the onset of neurological disease by later in life including dementia.

Neural MUA data was not significantly altered across any of the stimulations or conditions for any of the disease groups compared to WT controls. A consistent finding irrespective of stimulation and condition was that PCSK9-ATH mice display consistently reduced evoked-HbT responses (observed in chronic experiments) compared to WT controls, which suggests an advanced level of neurovascular breakdown and inefficiency. Other groups have found similarly reduced blood flow in the ApoE^-/-^model without altered cortical activation (Ayata et al., 2013). A recent study found decreased tissue oxygenation in the LDLR^-/-^mouse model of atherosclerosis (Li et al., 2019), and this is most likely to be the case in the PCSK9 model.

A question that arises is why the J20-PCSK9-MIX mice HbT responses are not more severely impaired than J20-AD and PCSK9-ATH? There may be redundancies that occur physiologically to compensate for mild hypoxia in the brain, such as the possible angiogenesis within the brain. Angiogenesis is known to be triggered in cerebral microvessels in AD in response to increased Aβ and neuroinflammation and may initially reflect a compensatory mechanism to increase perfusion (Jefferies et al., 2013). In addition, the levels of neuroinflammation seen in these mice may be due to an altered disease-course and examining temporospatial expression may reveal much higher levels of inflammation in this mixed model at an earlier time-point. Other markers of inflammation may be upregulated compared to those that we assessed, and future studies would incorporate transcriptomic approaches to identify other mechanisms or markers. Nevertheless, a key translational finding from our study was that J20-PCSK9-MIX mice displayed a significant increase in the number of hippocampal plaques.

There are several notable limitations with the present study. Firstly, all imaging was performed on lightly anaesthetised animals, which is known to compromise neurovascular function (Gao et al., 2017). However, previous research from our laboratory has developed an anaesthetic regimen that is comparable to awake imaging in terms of the haemodynamic responses to physiological whisker stimulation with little effect on vascular reactivity (Sharp et al., 2015). The benefits of lightly anaesthetised preparations over awake preparations is that we can avoid the multiple considerations of behavioural state in which the animals may be whisking, grooming as well as their arousal and stress states which may be present in awake animals. Furthermore, we report the stability and robustness of our imaging preparation in this study. We present the average of all the raw stimulation trials from each animal across the whole experimental session (Figure S2), showing the stability and robustness of our preparation, as well as easily identifying any changes. Secondly, our imaging analysis assumes O2 saturation to be 70% with a baseline haemoglobin concentration of 100μM. This may be important if the assumed baselines are different in the diseased animals compared to WT controls; however, our recent study (Sharp et al., 2019) using the same J20-AD mouse model discussed this issue in detail, in which we showed that regardless of the baseline blood volume estimation used, our percentage change was scaled by it (i.e. always the same change). Therefore, the observations in this paper with respect to the different diseased animals are robust.

In conclusion, we report novel findings of impaired neurovascular function in a novel experimental model of atherosclerosis (PCSK9-ATH) characterised by reduced stimulus-evoked blood volume without any significant alterations to evoked neural activity. We induced atherosclerosis in a mild fAD model (J20-AD) to create a mixed comorbid model (J20-PCSK9-ATH) in which we report a significant increase in the number of hippocampal Aβ plaques, however, without any significant changes to evoked haemodynamic or neural responses compared to WT or J20-AD mice. A key finding from this study was CSD was more severe in diseased animals. This may reflect the global inflammatory state of the brain and could also serve to be an effective preclinical and human clinical biomarker for baseline state and to assess therapies. Future studies should include assessment of other inflammatory markers and cellular pathway changes by a genome wide transcriptomics approach from single cell populations. It would also be prudent to induce atherosclerosis in a more severe fAD model to provide a severity continuum of mixed models that reflect clinical presentations of dementia.

## Materials & Methods

### Animals

All animal procedures were performed with approval from the UK Home Office in accordance to the guidelines and regulations of the Animal (Scientific Procedures) Act 1986 and were approved by the University of Sheffield ethical review and licensing committee. Male C57BL/6J mice were injected i.v at 6wks with 6×10^12^ virus molecules/ml rAAV8-mPCSK9-D377Y (Vector Core, Chapel Hill, NC) and fed a Western diet (21% fat, 0.15% cholesterol, 0.03% cholate, 0.296% sodium; #829100, Special Diet Services UK) for 8m (PCSK9-ATH). These mice were compared to age-matched wild-type C57BL/6J mice (with no AAV injection fed normal rodent chow) that were used as controls (WT C57BL/6J). In addition, male heterozygous transgenic J20-hAPP B6.Cg-Zbtb20Tg(PDGFB-APPSwInd)20Lms/2Mmjax) (MMRRC Stock No: 34836-JAX) mice were used. Atherosclerosis was induced in J20-hAPP mice alongside WT mice at 6wks of age combined with a Western diet to create a comorbid mixed model (J20-PCSK9-MIX). For the CSD imaging experiments, 4 nNOS-ChR2 mice (M/F, 16-40 weeks old) were included in the WT group. [nNOS-ChR2 mice: heterozygous nNOS-CreER (Jax 014541, (Taniguchi et al., 2011)) x homozygous Ai32 mice (Jax 024109, (Madisen et al., 2012)), given tamoxifen (100mg/kg, i.p., 3 injections over 5 days) at 1-2 months old]. All mice were imaged between 9-12m of age. All mice were housed in a 12hr dark/light cycle at a temperature of 23C, with food and water supplied *ad-libitum*.

### Thinned Cranial Window Surgery

Mice were anaesthetised with 7ml/kg i.p. injection of fentanyl-fluanisone (Hypnorm, Vetapharm Ltd), midazolam (Hypnovel, Roche Ltd) and maintained in a surgical anaesthetic plane by inhalation of isoflurane (0.6-0.8% in 1L/min O_2_). Core body temperature was maintained at 37°C through use of a homeothermic blanket (Harvard Apparatus) and rectal temperature monitoring. Mice were placed in a stereotaxic frame (Kopf Instruments, US) and the bone overlying the right somatosensory cortex was thinned forming a thinned cranial optical window. A thin layer of clear cyanoacrylate glue was applied over the cranial window to reinforce the window. Dental cement was applied around the window to which a metal head-plate was chronically attached. All mice were given 3 weeks to recover before the first imaging session.

### 2D-Optical Imaging Spectroscopy (2D-OIS)

2D-OIS measures changes in cortical haemodynamics: total haemoglobin (HbT), oxyhaemoglobin (HbO) and deoxyhaemoglobin (HbR) concentrations (Berwick et al., 2005). Mice were lightly sedated and placed into a stereotaxic frame. Sedation was induced as described above and maintained using low levels of isoflurane (0.3-0.6%). For imaging, the right somatosensory cortex was illuminated using 4 different wavelengths of light appropriate to the absorption profiles of the differing haemoglobin states (495nm ± 31, 559nm ± 16, 575nm ± 14 & 587nm ± 9) using a Lambda DG-4 high-speed galvanometer (Sutter Instrument Company, US). A Dalsa 1M60 CCD camera was used to capture the re-emitted light from the cortical surface. All spatial images recorded from the re-emitted light underwent spectral analysis based on the path length scaling algorithm (PLSA) as described previously (Berwick et al., 2005; Mayhew et al., 1999). which uses a modified Beer-Lambert law with a path light correction factor converting detected attenuation from the re-emitted light with a predicted absorption value. Relative HbT, HbR and HbO concentration estimates were generated from baseline values in which the concentration of haemoglobin in the tissue was assumed to be 100μM and O_2_ saturation to be 70%. For the stimulation experiments, whiskers were mechanically deflected for a 2s-duration and a 16s-duration at 5Hz using a plastic T-shaped stimulator which caused a 1cm deflection of the left-whisker. Each individual experiment consisted of 30 stimulation trials (for 2s) and 15 stimulation trials (for 16s) of which a mean trial was generated after spectral analysis of 2D-OIS. Stimulations were performed with the mouse breathing in 100% O_2_ or 21% O_2_, and a gas transition to medical air (21% O_2_) as well as an additional 10% CO_2_-hypercapnia test of vascular reactivity.

### Neural Electrophysiology

Simultaneous measures of neural activity alongside 2D-OIS were performed in a final acute imaging session 1-week after the 1st imaging session. A small burr-hole was drilled through the skull overlying the active region (as defined by the biggest HbT changes from 2D-OIS imaging) and a 16-channel microelectrode (100μm spacing, 1.5-2.7MΩ impedance, site area 177μm2) (NeuroNexus Technologies, USA) was inserted into the whisker barrel cortex to a depth of ~1500μm. The microelectrode was connected to a TDT preamplifier and a TDT data acquisition device (Medusa BioAmp/RZ5, TDT, USA). Multi-unit analysis (MUA) was performed on the data. All channels were depth aligned to ensure we had twelve electrodes covering the depth of the cortex in each animal. The data were high passed filtered above 300Hz to remove all low frequency components and split into 100ms temporal bins. Within each bin any data crossing a threshold of 1.5SD above the mean baseline was counted and the results presented in the form of fractional changes to MUA.

### Region Analysis

Analysis was performed using MATLAB (MathWorks). An automated region of interest (ROI) was selected using the stimulation data from spatial maps generated using 2D-OIS. The threshold for a pixel to be included within the ROI was set at 1.5xSD, therefore the automated ROI for each session per animal represents the area of the cortex with the largest haemodynamic response, as determined by the HbT. For each experiment, the response across all pixels within the ROI was averaged and used to generate a time-series of the haemodynamic response against time.

### Statistical Analysis

Statistical analyses were performed using SPSS v25 & GraphPad Prism v8. Shapiro-Wilks test was used to check for normality and Levene’s test was used to assess equality of variances. 2-way mixed design ANOVA, 1-way ANOVA or Kruskal-Wallis tests were used, as appropriate. For 1-way ANOVA, if variances were unequal, Welch’s F was reported. Results were considered statistically significant if p<0.05. The Shapiro-Wilks test suggested that, for chronic experiments, peak values of HbT and HbO are normally distributed, however, HbR values are significantly non-normal. 2-way mixed design was used to compare peak values for HbT, HbO & HbR (although HbR failed the S-W test for normality, an ANOVA was used as they were considered fairly robust against small deviations from normality). Inspection of Levene’s test suggested that variances were equal, therefore, Dunnett’s (two-sided) multiple comparisons test was used to compare disease models to WT, and for HbR, Games-Howell multiples comparisons were used. If the Greenhouse-Geisser estimate of sphericity showed deviation from sphericity (chronic experiments: HbT (ε=0.55), HbO (ε = 0.49) & HbR (ε=0.564), results are reported with Greenhouse-Geisser correction applied. qRT-PCR data was analysed by performing 1-way ANOVAs with Dunnett’s multiple comparisons test used to compare disease models to WT. P-values <0.05 were considered statistically significant. All the data are presented as mean values ± standard error of mean (SEM).

### Immunohistochemistry

At the end of terminal experiments, mice were euthanized with an overdose of pentobarbital (100mg/kg, Euthatal, Merial Animal Health Ltd) and transcardially perfused with 0.9% saline and brains were dissected. One half-hemisphere of the brains were fixed in formalin and embedded in paraffin wax, with the other half snap-frozen using isopentane and stored at −80C. 5μm coronal sections were obtained using a cryostat. Immunohistochemistry was performed using an avidin-biotin complex (ABC) method (as described previously (Ameen-Ali et al., 2019)). Following slide preparation and antigen retrieval (pressure cooker at 20psi at 120C for 45s (pH6.5)), sections underwent additional pre-treatment in 70% formic acid. Sections were incubated with 1.5% normal serum followed by incubation with the primary antibody (biotinylated anti-Aβ – 1:100, BioLegend, USA) for 1 hour. Horseradish peroxidase avidin-biotin complex (Vectastain Elite Kit, Vector Laboratories, UK) was used to visualise antibody binding along with 3,3-diaminobenzidine tetrahydrochloride (DAB) (Vector Laboratories, UK). All sections were counterstained with haematoxylin, dehydrated and mounted in DPX. Sections were imaged using a Nikon Eclipse Ni-U microscope attached to a Nikon DS-Ri1 camera. Plaques were identified at x40 magnification and manually counted per section.

### qRT-PCR

Snap-frozen hemispheres were homogenised, and RNA was extracted using Direct-zol RNA MiniPrep kit with TRI-reagent as per the manufacturer’s guidelines (Zymo) and RNA quality checked using NanoDropTM (ThermoFisher Scientific). cDNA was synthesised from the extracted RNA using the UltraScript 2.0 cDNA synthesis kit (PCR BioSystems) according to the manufacturer’s guidelines. qRT-PCR was performed using PrimeTime qRT-PCR assay primers (IDT) for *IL1β* & *TNFα* with *ACTB* as the reference housekeeping gene. Luna qRT-PCR Master Mix (NEB) was used with the primers, cDNA and nuclease free water and each gene for each sample was duplicated. CFX384 Real-Time System (BioRad) with a C1000 Touch Thermal Cycler (BioRad) was used to perform qRT-PCR consisting of 40 cycles. Data was analysed using the well-established delta-Ct method (Livak & Schmittgen, 2001) by normalising against *ACTB*.

---

**Figure Supplemental 1:**
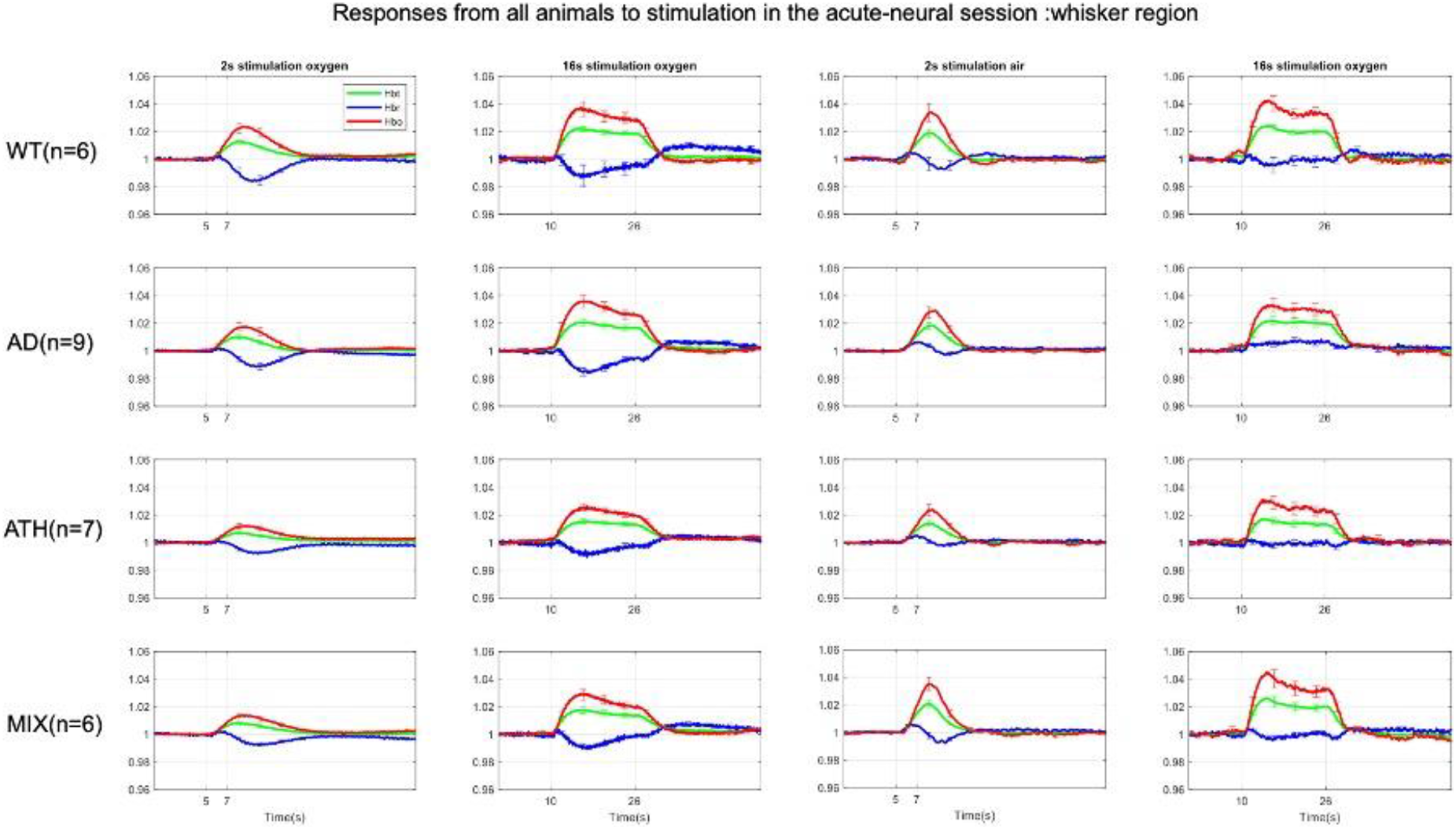
Fractional Changes in Acute Stimulus-Evoked Haemodynamic Responses. HbT: There was no significant overall effect of disease F(3,24)=1.69, p=0.196. As expected, there was a significant effect of experiment, F(2.16,51.73)=76.72, p<0.001. There was no significant interaction effect between experiment and disease, F(6.47,51.73)=1.73, p=0.127. **HbO:** There was no significant overall effect of disease F(3,24)=1.36, p=0.280. There was a significant effect of experiment, F(2.02,48.57=62.10, p<0.001. There was no significant interaction effect between experiment and disease, F(6.07,48.57)=2.08, p=0.072. **HbR:** There was no significant overall effect of disease F(3,24)=1.42, p=0.262. As expected, there was a significant effect of experiment, F(2.18,52.42)=17.54, p<0.001. There was a significant interaction effect between experiment and disease F(6.55, 52.42)=2.54, p=0.028. All error bars (lightly shaded) are ±SEM. Vertical dotted lines indicate start and end of stimulations.

**Figure Supplemental 2:**
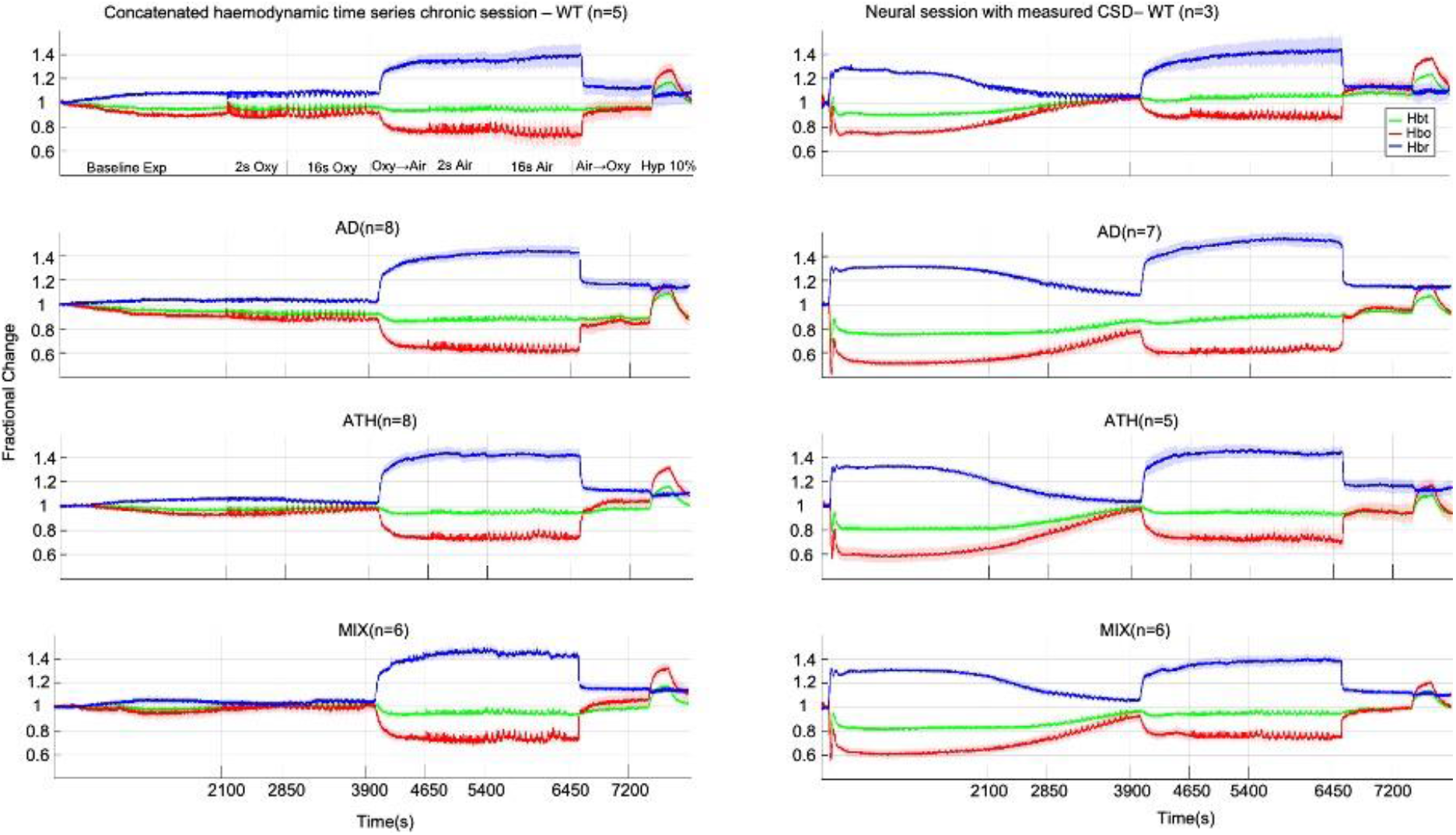
Concatenated data showing stability and robustness of the mouse imaging preparation. Left) Chronic imaging sessions including a 35-minute haemodynamics baseline before first stimulation. Right) Acute imaging sessions including CSD plus 35-minute haemodynamics recovery before first stimulation.

**Figure Supplemental 3:**
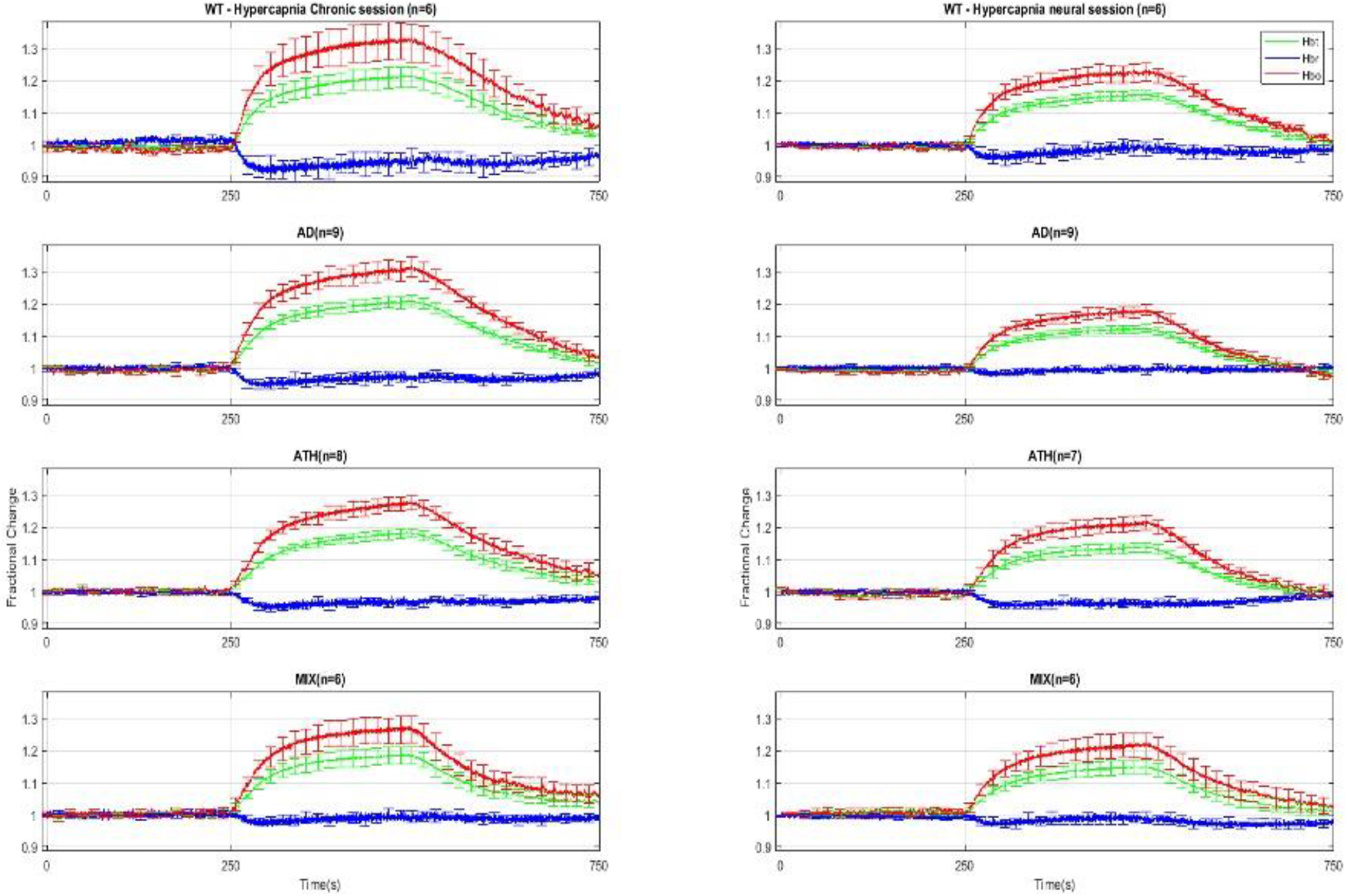
Chronic (Left) & Acute (Right) Hypercapnia. Chronic: A 1-way ANOVA showed no significant effect of disease for HbT (F(3,12.06)=0.49, p=0.694), HbO (F(3,11.98)=0.44, p=0.732) nor HbR (F(3,12.081)=0.98, p =0.436). Acute: A 1-way ANOVA showed no significant effect of disease for HbO F(3,12.00)=0.74, p=0.549 but there was a significant effect of disease for HbR F(3,11.01)=3.77, p=0.044. Games-Howell multiple comparisons showed that, for HbR, there was a significant difference between AD and ATH (p=0.019). Kruskal-Wallis test showed no significant effect of disease for HbT: H(3)=2.87, p=0.412. Error bars ±SEM.

## Acknowledgments

OS’s PhD studentship and consumables were funded by the Neuroimaging in Cardiovascular Disease (NICAD) network scholarship (University of Sheffield). A British Heart Foundation (BHF) project grant was awarded to SEF to carry out the work using the PCSK9 model (PG/13/55/30365). The J20-mouse colony was in part funded and supported by Alzheimer’s Research UK (Grant R/153749-12-1). CH is funded by a Sir Henry Dale Fellowship jointly funded by the Wellcome Trust and the Royal Society. This research was funded in whole, or in part, by the Wellcome Trust [Grant number 105586/Z/14/Z]. For the purpose of Open Access, the author has applied a CC BY public copyright licence to any Author Accepted Manuscript version arising from this submission. MAR is funded by a Conacyt scholarship. We would like to thank Prof Lennart Mucke (Gladstone Institute of Neurological Disease & Department of Neurology, UCSF, CA, US) as well as the J. David Gladstone Institutes for the J20-hAPP mice. Finally, we would like to thank Mr Michael Port for building and maintaining the imaging apparatus and Dr Luke Boorman for producing MATLAB code for data analysis.

## Author Contributions

OS performed the majority of the *in vivo* experiments and authored the manuscript. OS & JB designed the experiments. OS, BP, LL, BE, PS & MAR performed experiments. OS, CH & JB performed MATLAB and data analysis. JB, SEF, CH, PRH & SBW supervised the research and provided editorial guidance. All authors proofread the final version of the manuscript.

## References

Ameen-Ali, K. E., Simpson, J. E., Wharton, S. B., Heath, P. R., Sharp, P. S., Brezzo, G., & Berwick, J. (2019). The Time Course of Recognition Memory Impairment and Glial Pathology in the hAPP-J20 Mouse Model of Alzheimer’s Disease. J Alzheimers Dis, 68(2), 609–624. https://doi.org/10.3233/JAD-181238

Ayata, C., & Lauritzen, M. (2015). Spreading Depression, Spreading Depolarizations, and the Cerebral Vasculature. Physiol Rev, 95(3), 953–993. https://doi.org/10.1152/physrev.00027.2014

Ayata, C., Shin, H. K., Dilekoz, E., Atochin, D. N., Kashiwagi, S., Eikermann-Haerter, K., & Huang, P. L. (2013). Hyperlipidemia disrupts cerebrovascular reflexes and worsens ischemic perfusion defect. J Cereb Blood Flow Metab, 33(6), 954–962. https://doi.org/10.1038/jcbfm.2013.38

Berwick, J., Johnston, D., Jones, M., Martindale, J., Redgrave, P., McLoughlin, N., Schiessl, I., & Mayhew, J. E. (2005). Neurovascular coupling investigated with two-dimensional optical imaging spectroscopy in rat whisker barrel cortex. Eur J Neurosci, 22(7), 1655–1666. https://doi.org/10.1111/j.1460-9568.2005.04347.x

Bjorklund, M. M., Hollensen, A. K., Hagensen, M. K., Dagnaes-Hansen, F., Christoffersen, C., Mikkelsen, J. G., & Bentzon, J. F. (2014). Induction of atherosclerosis in mice and hamsters without germline genetic engineering. Circ Res, 114(11), 1684–1689. https://doi.org/10.1161/CIRCRESAHA.114.302937

Dreier, J. P. (2011). The role of spreading depression, spreading depolarization and spreading ischemia in neurological disease. Nat Med, 17(4), 439–447. https://doi.org/10.1038/nm.2333

Eikermann-Haerter, K., Yuzawa, I., Dilekoz, E., Joutel, A., Moskowitz, M. A., & Ayata, C. (2011). Cerebral autosomal dominant arteriopathy with subcortical infarcts and leukoencephalopathy syndrome mutations increase susceptibility to spreading depression. Ann Neurol, 69(2), 413–418. https://doi.org/10.1002/ana.22281

Gao, Y. R., Ma, Y., Zhang, Q., Winder, A. T., Liang, Z., Antinori, L., Drew, P. J., & Zhang, N. (2017). Time to wake up: Studying neurovascular coupling and brain-wide circuit function in the un-anesthetized animal. Neuroimage, 153, 382–398. https://doi.org/10.1016/j.neuroimage.2016.11.069

Iadecola, C. (2017). The Neurovascular Unit Coming of Age: A Journey through Neurovascular Coupling in Health and Disease. Neuron, 96(1), 17–42. https://doi.org/10.1016/j.neuron.2017.07.030

Iadecola, C., Duering, M., Hachinski, V., Joutel, A., Pendlebury, S. T., Schneider, J. A., & Dichgans, M. (2019). Vascular Cognitive Impairment and Dementia: JACC Scientific Expert Panel. J Am Coll Cardiol, 73(25), 3326–3344. https://doi.org/10.1016/j.jacc.2019.04.034

Jefferies, W. A., Price, K. A., Biron, K. E., Fenninger, F., Pfeifer, C. G., & Dickstein, D. L. (2013). Adjusting the compass: new insights into the role of angiogenesis in Alzheimer’s disease. Alzheimers Res Ther, 5(6), 64. https://doi.org/10.1186/alzrt230

Kapasi, A., DeCarli, C., & Schneider, J. A. (2017). Impact of multiple pathologies on the threshold for clinically overt dementia. Acta Neuropathol, 134(2), 171–186. https://doi.org/10.1007/s00401-017-1717-7

Kapasi, A., & Schneider, J. A. (2016). Vascular contributions to cognitive impairment, clinical Alzheimer’s disease, and dementia in older persons. Biochim Biophys Acta, 1862(5), 878–886. https://doi.org/10.1016/j.bbadis.2015.12.023

Kurth, T., Rist, P. M., Ridker, P. M., Kotler, G., Bubes, V., & Buring, J. E. (2020). Association of Migraine With Aura and Other Risk Factors With Incident Cardiovascular Disease in Women. JAMA, 323(22), 2281–2289. https://doi.org/10.1001/jama.2020.7172

Li, B., Lu, X., Moeini, M., Sakadzic, S., Thorin, E., & Lesage, F. (2019). Atherosclerosis is associated with a decrease in cerebral microvascular blood flow and tissue oxygenation. PLoS One, 14(8), e0221547. https://doi.org/10.1371/journal.pone.0221547

Livak, K. J., & Schmittgen, T. D. (2001). Analysis of relative gene expression data using real-time quantitative PCR and the 2(-Delta Delta C(T)) Method. Methods, 25(4), 402–408. https://doi.org/10.1006/meth.2001.1262

Lusis, A. J. (2000). Atherosclerosis. Nature, 407(6801), 233–241. https://doi.org/10.1038/35025203

Madisen, L., Mao, T., Koch, H., Zhuo, J. M., Berenyi, A., Fujisawa, S., Hsu, Y. W., Garcia, A. J., 3rd, Gu, X., Zanella, S., Kidney, J., Gu, H., Mao, Y., Hooks, B. M., Boyden, E. S., Buzsaki, G., Ramirez, J. M., Jones, A. R., Svoboda, K., Han, X., Turner, E. E., & Zeng, H. (2012). A toolbox of Cre-dependent optogenetic transgenic mice for light-induced activation and silencing. Nat Neurosci, 15(5), 793–802. https://doi.org/10.1038/nn.3078

Matthews, F. E., Brayne, C., Lowe, J., McKeith, I., Wharton, S. B., & Ince, P. (2009). Epidemiological pathology of dementia: attributable-risks at death in the Medical Research Council Cognitive Function and Ageing Study. PLoS Med, 6(11), e1000180. https://doi.org/10.1371/journal.pmed.1000180

Mayhew, J., Zheng, Y., Hou, Y., Vuksanovic, B., Berwick, J., Askew, S., & Coffey, P. (1999). Spectroscopic analysis of changes in remitted illumination: the response to increased neural activity in brain. Neuroimage, 10(3 Pt 1), 304–326. https://doi.org/10.1006/nimg.1999.0460

Morton, R. E., St John, P. D., & Tyas, S. L. (2019). Migraine and the risk of all-cause dementia, Alzheimer’s disease, and vascular dementia: A prospective cohort study in community-dwelling older adults. Int J Geriatr Psychiatry, 34(11), 1667–1676. https://doi.org/10.1002/gps.5180

Mucke, L., Masliah, E., Yu, G. Q., Mallory, M., Rockenstein, E. M., Tatsuno, G., Hu, K., Kholodenko, D., Johnson-Wood, K., & McConlogue, L. (2000). High-level neuronal expression of abeta 1-42 in wild-type human amyloid protein precursor transgenic mice: synaptotoxicity without plaque formation. J Neurosci, 20(11), 4050–4058.

Munting, L. P., Derieppe, M., Suidgeest, E., Hirschler, L., van Osch, M. J., Denis de Senneville, B., & van der Weerd, L. (2021). Cerebral blood flow and cerebrovascular reactivity are preserved in a mouse model of cerebral microvascular amyloidosis. Elife, 10. https://doi.org/10.7554/eLife.61279

Napoli, C., Witztum, J. L., de Nigris, F., Palumbo, G., D’Armiento, F. P., & Palinski, W. (1999). Intracranial arteries of human fetuses are more resistant to hypercholesterolemia-induced fatty streak formation than extracranial arteries. Circulation, 99(15), 2003–2010. https://doi.org/10.1161/01.cir.99.15.2003

Neuropathology Group. Medical Research Council Cognitive, F., & Aging, S. (2001). Pathological correlates of late-onset dementia in a multicentre, community-based population in England and Wales. Neuropathology Group of the Medical Research Council Cognitive Function and Ageing Study (MRC CFAS). Lancet, 357(9251), 169–175. https://doi.org/10.1016/s0140-6736(00)03589-3

Rahimi, J., & Kovacs, G. G. (2014). Prevalence of mixed pathologies in the aging brain. Alzheimers Res Ther, 6(9), 82. https://doi.org/10.1186/s13195-014-0082-1

Ripa, P., Ornello, R., Pistoia, F., Carolei, A., & Sacco, S. (2015). Spreading depolarization may link migraine, stroke, and other cardiovascular disease. Headache, 55(1), 180–182. https://doi.org/10.1111/head.12436

Roche-Molina, M., Sanz-Rosa, D., Cruz, F. M., Garcia-Prieto, J., Lopez, S., Abia, R., Muriana, F. J., Fuster, V., Ibanez, B., & Bernal, J. A. (2015). Induction of sustained hypercholesterolemia by single adeno-associated virus-mediated gene transfer of mutant hPCSK9. Arterioscler Thromb Vasc Biol, 35(1), 50–59. https://doi.org/10.1161/ATVBAHA.114.303617

Shabir, O., Berwick, J., & Francis, S. E. (2018). Neurovascular dysfunction in vascular dementia, Alzheimer’s and atherosclerosis. BMC Neurosci, 19(1), 62. https://doi.org/10.1186/s12868-018-0465-5

Shabir, O., Sharp, P., Rebollar, M. A., Boorman, L., Howarth, C., Wharton, S. B., Francis, S. E., & Berwick, J. (2020). Enhanced Cerebral Blood Volume under Normobaric Hyperoxia in the J20-hAPP Mouse Model of Alzheimer’s Disease. Sci Rep, 10(1), 7518. https://doi.org/10.1038/s41598-020-64334-4

Sharp, P. S., Ameen-Ali, K. E., Boorman, L., Harris, S., Wharton, S., Howarth, C., Shabir, O., Redgrave, P., & Berwick, J. (2019). Neurovascular coupling preserved in a chronic mouse model of Alzheimer’s disease: Methodology is critical. J Cereb Blood Flow Metab, 271678X19890830. https://doi.org/10.1177/0271678X19890830

Sharp, P. S., Shaw, K., Boorman, L., Harris, S., Kennerley, A. J., Azzouz, M., & Berwick, J. (2015). Comparison of stimulus-evoked cerebral hemodynamics in the awake mouse and under a novel anesthetic regime. Sci Rep, 5, 12621. https://doi.org/10.1038/srep12621

Taniguchi, H., He, M., Wu, P., Kim, S., Paik, R., Sugino, K., Kvitsiani, D., Fu, Y., Lu, J., Lin, Y., Miyoshi, G., Shima, Y., Fishell, G., Nelson, S. B., & Huang, Z. J. (2011). A resource of Cre driver lines for genetic targeting of GABAergic neurons in cerebral cortex. Neuron, 71(6), 995–1013. https://doi.org/10.1016/j.neuron.2011.07.026

Yemisci, M., & Eikermann-Haerter, K. (2019). Aura and Stroke: relationship and what we have learnt from preclinical models. J Headache Pain, 20(1), 63. https://doi.org/10.1186/s10194-019-1016-x

Zlokovic, B. V. (2011). Neurovascular pathways to neurodegeneration in Alzheimer’s disease and other disorders. Nat Rev Neurosci, 12(12), 723–738. https://doi.org/10.1038/nrn3114

